# New intranasal and injectable gene therapy for healthy life extension

**DOI:** 10.1101/2021.06.26.449305

**Authors:** Dabbu Kumar Jaijyan, Anca Selariu, Ruth Cruz-Cosme, Mingming Tong, Shaomin Yang, George Church, David Kekich, Ali Fallah, Junichi Sadoshima, Qiyi Tang, Elizabeth Parrish, Hua Zhu

**Author notes:** Co-first authors, these authors contributed equally to this work (Dabbu Kumar Jaijyan, Anca Selariu). Correspondence and requests for materials should be addressed to: Dr. Hua Zhu, Department of Microbiology, Biochemistry and Molecular Genetics, New Jersey Medical School, Rutgers the State University of New Jersey, 225 Warren Street, Newark, New Jersey, 07103, USA;, Elizabeth Parrish, Chief Executive Officer, BioViva USA Inc, Delaware, United States.

## Abstract

As the global elderly population grows, it is socioeconomically and medically critical to have diverse and effective means of mitigating the impact of aging on human health. Previous studies showed that adenovirus-associated virus (AAV) vector induced overexpression of certain proteins can suppress or reverse the effects of aging in animal models. Here, we sought to determine whether the high-capacity cytomegalovirus vector can be an effective and safe gene delivery method for two such-protective factors: telomerase reverse transcriptase (TERT) and follistatin (FST). We found that the mouse cytomegalovirus (MCMV) carrying exogenous TERT or FST (MCMV_TERT_ or MCMV_FST_) extended median lifespan by 41.4% and 32.5%, respectively. This is the first report of CMV being used successfully as both an intranasal and injectable gene therapy system to extend longevity. Treatment significantly improved glucose tolerance, physical performance, and prevented loss of body mass and alopecia. Telomere shortening seen with aging was ameliorated by TERT, and mitochondrial structure deterioration was halted in both treatments. Intranasal and injectable preparations performed equally well in safely and efficiently delivering gene therapy to multiple organs, with long-lasting benefits and without carcinogenicity or unwanted side effects. Translating this research to humans could have significant benefits associated with increased health span.

## Introduction

How to achieve healthy longevity has remained a challenging subject in biomedical science. It has been well established that aging is associated with a reduction in telomere repeat elements at the ends of chromosomes (1), which in part results from insufficient telomerase activity. Importantly, the biological functions of the telomerase complex rely on telomerase reverse transcriptase (TERT) (2). TERT plays a major role in telomerase activation, and telomerase lengthens the telomere DNA (2, 3). Because telomerase supports cell proliferation and division by reducing the erosion of chromosomal ends (4) in mitotic cells, animals deficient in TERT have shorter telomeres and shorter life spans(5, 6). Recent studies on animal models have shown the therapeutic efficacy of TERT in increasing healthy longevity and reversing the aging process (7–9). Telomere shortening also increases the risk of heart disease by a mechanism that remains unclear (10). The follistatin (FST) gene encodes a monomeric secretory protein that is expressed in nearly all mammalian tissues. In muscle cells, FST functions as a negative regulator of myostatin, a myogenesis inhibitory signal protein (11). FST overexpression is known to increase skeletal muscle mass in transgenic mice by 194-327%(12) by neutralizing the effects of various TGF-β ligands involved in muscle fiber break-down, including myostatin and activin inhibition complex (13). FST- knockout mice have smaller and fewer muscle fibers, and show retarded growth, skeletal defects, reduced body mass, and die in a few hours after birth, suggesting an important role of FST in skeletal muscle development (11, 14). These findings strongly implicate the therapeutic potential of FST in the treatment of muscular dystrophy and muscle loss caused by aging or microgravity. Thus, TERT and FST are among prime candidates for gene therapy aimed to improve healthy life spans.

As more longevity-supporting factors are discovered, it is of interest to determine potential large capacity vectors for delivering multiple genes simultaneously. Unlike AAV, lentiviruses or other viral vectors used for gene delivery, cytomegaloviruses have a large genome size and unique ability to incorporate multiple genes. Cytomegaloviruses also do not integrate their DNA into the host genome during the infection cycle, thus mitigating the risk of insertional mutagenesis. They also do not elicit symptomatic immune reactions in most healthy hosts (15). Notably, the CMV vector does not invoke genome instability and has not been identified to cause malignancies (16, 17). Human CMV (HCMV) has been proven a safe delivery vector for expressing therapeutic proteins in human clinical trials (16). MCMV and HCMV are similar in many aspects, including viral pathogenesis, homology, viral protein function, viral gene expression and viral replication. Using mouse cytomegalovirus (MCMV) as a viral vector, we examined the therapeutic potential of TERT and FST gene therapy to offset biological aging in a mouse model, and demonstrated significant lifespan increase, as well as positive metabolic and physical performance effects. Further studies may elucidate the full CMV cargo capacity and effectiveness. Translational studies are required to determine whether our findings can be replicated in human subjects.

## Materials and methods

### Construction of MCMV with a luciferase marker

A firefly luciferase expression cassette was inserted into MCMV genome by replacing a nonessential gene *ie2* using the BACmid Sm3fr (Smith strain) (18). The seamless BAC system using *galK* as a selection marker (19, 20) was utilized to construct the luciferase expressing MCMV_Luc_. First, the *ie2* gene was replaced with the *galK* gene by PCR using primers Fw: ccccctccggggcgagtcttttacaggctacaacgactgtccgatgaataCCTGTTGACAATTAATCATCGGCA and Rv: catcccgggagggccagcgtagtctccgttgctgggttcggccgagggtTCAGCACTGTCCTGCTCCTT; then, the *galK* gene was replaced by a luciferase expression cassette which was amplified by PCR using primers Fw: ccccctccggggcgagtcttttacaggctacaacgactgtccgatgaataGATATACGCGTTGACATTGA and Rv: catcccgggagggccagcgtagtctccgttgctgggttcggccgagggatTCAGACAATGCGATGCAATTTC. The BAC DNA was transfected into NIH/3T3 cells to generate recombinant virus. The resultant MCMV is named as MCMV_Luc_. (referred to as WT in this study) and its growth kinetics and luciferase expression were analyzed using standard methods (21). Replication was observed and measured in live animals using the In Vivo Imaging System (IVIS).

### Construction and characterization of recombinant MCMV vectors

A BAC engineering method was used to generate recombinant MCMV_TERT_ and MCMV_FST_ containing mouse telomerase reverse transcriptase (TERT) and mouse follistatin (FST) expression cassettes, respectively, as described previously (22). The TERT and FST expression cassettes containing a FLAG-tag at the C-terminus were cloned from the plasmids, MR226892 and MR225488 (Origene). The TERT and FST expression cassettes were inserted between the M106 and M107 locus of MCMV_Luc_ (WT) genome without any deletion to generate MCMV_TERT_ and MCMV_FST_ viruses. Expression was controlled by the HCMV major IE promoter. Virus preparation and growth curve analysis were performed as described previously (23, 24). For the western blot analysis, NIH/3T3 cells were infected with MCMV_TERT_ and MCMV_FST_ and western blot was performed using mouse anti-FLAG antibody.

### Measurement of telomere length

The heart, brain, liver, kidney, lung, and muscle were harvested from MCMV_TERT_, MCMV_FST_, WT, and untreated mice at the age of 24 months. An 8-month-old mouse was used as a control. The tissues were homogenized, and genomic DNA was isolated using Sigma genomic DNA isolation kit (GeneElute genomic DNA isolation kit). The real-time PCR was performed in the CFX96 real-time PCR system (Bio-Rad), as described previously (25, 26), using specific primers for telomere, forward, and reverse telomeric primers are 5’ CGG TTT GTT TGG GTT TGG GTT TGG GTT TGG GTT TGG GTT 3’ and 5’ GGC TTG CCT TAC CCT TAC CCT TAC CCT TAC CCT TAC CCT 3’ respectively (25). A single copy conserved gene, the acidic ribosomal phosphoprotein (36B4) gene, was used as an internal control. Forward and reverse primers for 36B4 were 5’ ACT GGT CTA GGA CCCGAG AAG 3’ and 5’ TCA ATG GTG CCT CTG GAG ATT 3’ respectively (25). The relative telomere length was calculated by ΔCT value as described previously (26).

### Measurement of TERT and FST expression in different tissue

Eleven-month-old C57BL/6J female mice (3 per group) were treated with WT, MCMV_TERT_ and MCMV_FST_. The heart, brain, lung, liver, muscle, and kidneys were isolated 6 days post-inoculation. RNA was extracted using RNeasy^®^ Mini Kit (Qiagen). Approximately, 1.0μg of total RNA was used to prepare cDNA using Titanium RT-PCR kit (TaKaRa). The real-time PCR was performed on the cDNA using mouse TERT and FST primers as described previously (27, 28). β-actin was used as an internal control (29).

### Determination of TERT and FST protein levels in the sera

Preliminary TERT protein expression kinetics was performed using mock, WT and MCMV_TERT_ treated 8-month-old C57BL/6J mice via both IN and IP routes. Blood samples were collected at day 0, 3, 5, 7, 9, 12, 15, 20, 25 and 30 post inoculation, and sera were prepared using ELISA kits MBS1601022 and MBS1996306 respectively (MyBioSource). For the longevity study, blood was collected from 24-month-old treated mice from each group, and sera were analyzed 4 days post treatment to determine the level of mouse TERT and FST in blood sera.

### Bodyweight and body hair analyses

The bodyweights and body hair loss of all mice in each group were measured and recorded twice a month until mice died. Representative photographs of coat and fur characteristics were taken during the 6th months of treatment.

### Activity test and beam coordination test

Three 24-month-old mice from each group were used in these tests. For the beaker escape test (24, 30), mice were placed individually in a 1L glass beaker and the number of times each mouse tried to climb on the wall of the beaker was recorded. For the beam coordination test (31), mice were trained to traverse a 4-foot-long and 1-cm-wide beam for two consecutive days, then tested on the third day. The time required by each mouse to cross the beam was measured.

### Glucose tolerance and glycosylated hemoglobin A1c (HbA1c) test

Three 22-month-old mice from each group were subjected for a glucose tolerance test as described previously (32). Briefly, the mice were starved for 15 hours, then injected intraperitoneally with 50mg glucose. Blood samples were collected at 0, 15, 30, 60, 120, 180, 240, 300, 360, 420 and 480 minutes, and the blood glucose levels were immediately determined using OneTouch Ultra glucose meter. For the measurement of HbA1c, 100μl blood samples were collected from three 23-month-old mice in each group. The blood samples were allowed to clot at RT, and sera were prepared. The sera were analyzed by the ELISA kit (80310, Crystal Chem).

### Transmission electron microscopy (TEM)

Two mice from each treated group (one from IP, one from IN) were sacrificed at 24 months. The heart and skeletal muscle were isolated for EM analysis. The tissue samples were fixed in EM buffer (Electron Microscopic Sciences). TEM and mitochondrial analysis were performed as described previously (33).

### Statistical analyses

The data was analyzed by unpaired t-test using R environment (version 3.4.5) with ggplot software and a p-value <0.05 was considered significant. The survival curve of mice in each group was determined by Kaplan-Meier survival curve.

## Results

### Construction of MCMV_TERT_ and MCMV_FST_

We developed an MCMV vector that expresses luciferase as a reporter gene (MCMV_Luc_) to easily monitor MCMV infection and cellular replication in cell culture and in a mouse model. MCMV_Luc_ replicated as well as its parental virus, and we used it throughout this study as empty viral vector control or wild-type MCMV virus inoculation (WT) (21).

We constructed recombinant MCMV vectors expressing FLAG-tagged genes TERT and FST genes (MCMV_TERT_ and MCMV_FST_) and demonstrated that they replicated as productively as MCMV_Luc_ (WT) in mouse fibroblast cells and *in vivo* (Fig. 1A-D).

**Fig. 1.**
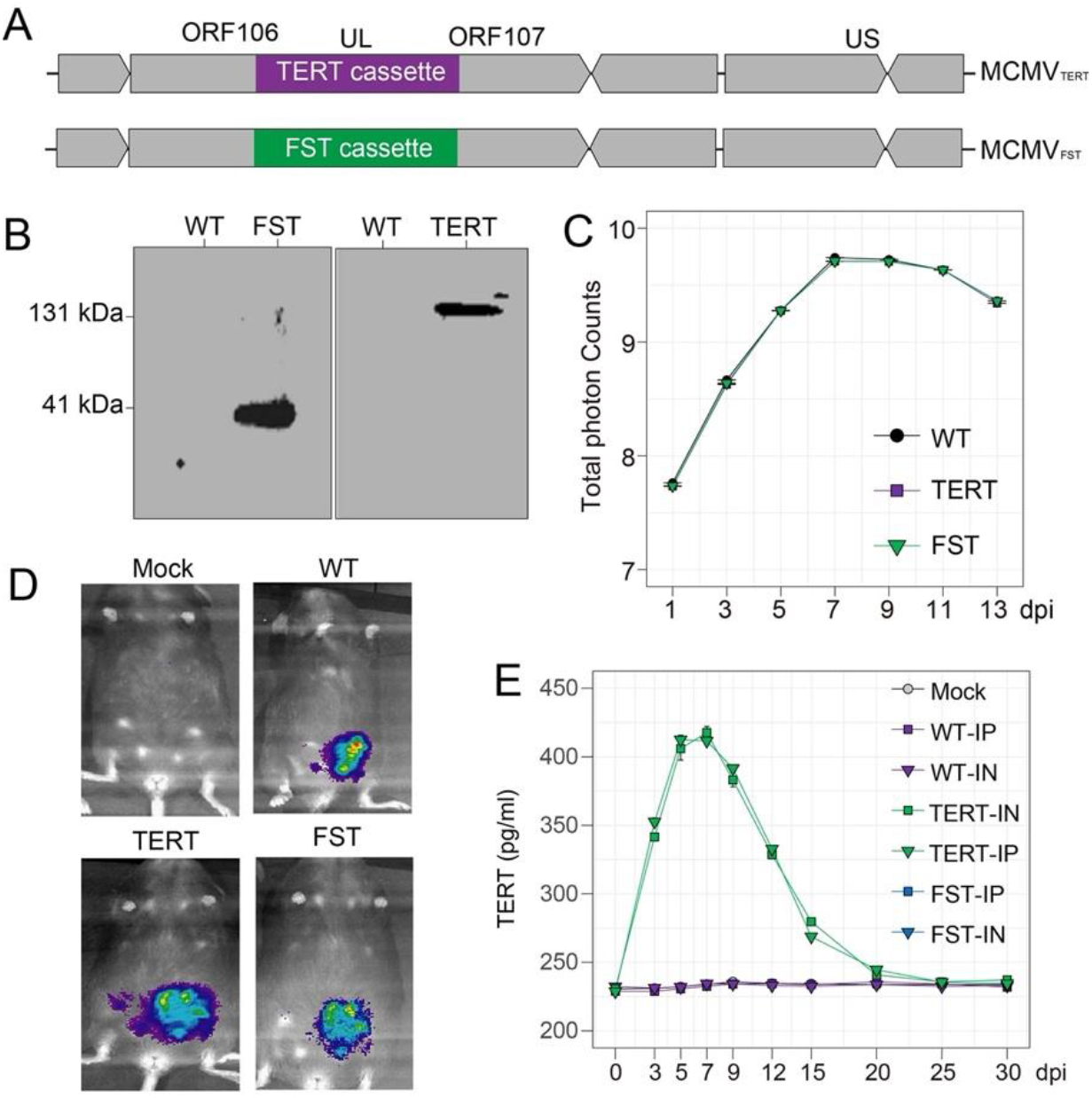
Construction and verification of MCMV_TERT_ and MCMV_FST_. (**A**) TERT-3’ FST-3’ FLAG constructs. (**B**) Expression of TERT (131 kDa) or FST (41 kDa) proteins in MCMV_TERT_ or MCMV_FST_ treated NIH/3T3 cells. (**C**) Plaque formation unit (PFU) assay growth curve of MCMV_LUC_ (WT), MCMV_TERT_, and MCMV_FST_ in NIH/3T3. (**D**) Luciferase signal *in vivo* 3 days after IP inoculations with mock, WT, MCMV_TERT_, and MCMV_FST_. (**E**) Detection of TERT by ELISA in serum of MCMV_TERT_ treated 8-month-old mice over one month. Each data point represents the average value of TERT in three mice.. Error bars represent the standard deviations.

TERT protein expression delivered Intraperitoneally (IP) or intranasally (IN) peaked at 7 days, and then gradually decreased, reaching the basal level at around day 25 (Fig. 1E), confirming the vector’s ability to deliver exogenous proteins *in vivo.* This was further confirmed by the analysis of TERT and FST mRNAs and proteins in blood and tissues harvested from treated animals, as described below.

### Significant lifespan extension

Seven groups of eight aged female C57BL/6J mice received mock (IP), WT-IN, WT-IP, MCMV_TERT_-IN, MCMV_TERT_-IP, MCMV_FST_-IN, and MCMV_FST_ -IP, respectively, for six consecutive months, at doses of 1×10^5^ PFU. Treatment started in 18-month-old mice, equivalent to approximately 56-years-old in humans (Fig. 2B) (34). One mouse per group was sacrificed at 24 months for tissue analyses, while the remaining subjects were monitored for physical and physiological changes until their natural death.

**Fig. 2.**
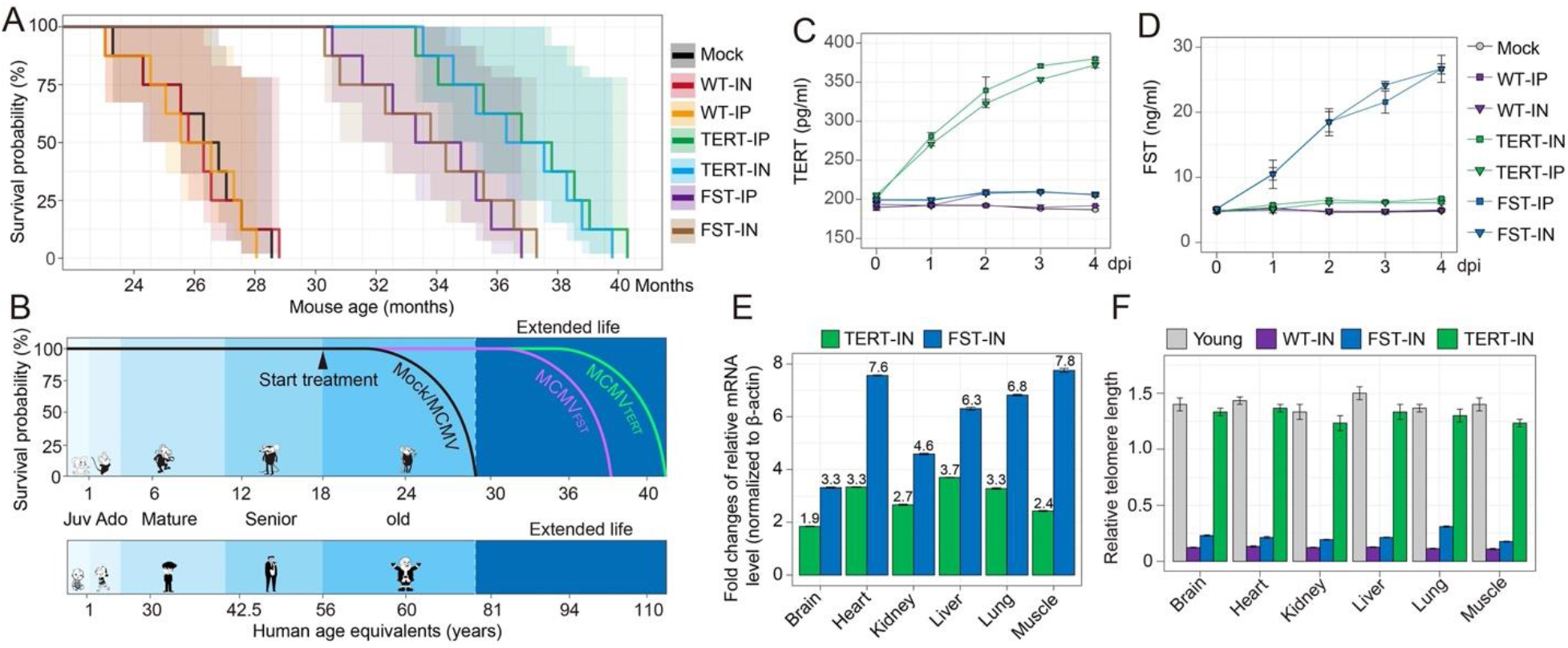
MCMV_TERT_ and MCMV_FST_ significantly extend lifespan. (**A**) Survivorship curve comparison, 8 mice per group. (**B**) C57BL/6J mice and human age equivalence at the start of experimental treatment (black arrow; adapted from Fox JG et al. (34)). (**C**) TERT and (**D**) FST proteins by ELISA in blood cell extracts from 24-month-old mice.. Error bars represent the standard deviations. (**E**) The fold increase of TERT and FST mRNA levels in organs of WT, MCMV_TERT_, and MCMV_FST_ treated mice by qPCR. (**F**) Relative telomere length in organs of 24-month-old mice vs an 8-month-old control. 36B4 gene was used for normalization.

Mock and WT controls died at 26.7, 26.5, and 26.4 months (median), consistent with previous reports on lifespan of female C57BL/6J mice (35, 36). The median age at death in MCMV_FST_ treated groups was 35.1 months (32.5% increase), while MCMV_TERT_ treated mice lived 37.5 months (41.4% increase) (Fig. 2A and B, Supplementary Table 1S). This result exceeds the longevity achieved with a single dose of AVV9-TERT in the same animal model (13-24% when delivered in a single dose at 2 and 1 year old mice, respectively) (7). Interestingly, CMV therapy was equally effective regardless of route of inoculation, although the mechanism of dissemination differs, suggesting that expression of the therapeutic load is not substantially affected by the vector’s interaction with the immune system (37) (38).

### Systemic TERT and FST expression

The amounts of TERT (Fig. 2C) or FST (Fig. 2D) proteins in blood increased daily in the first four days post-inoculation, while endogenous protein levels remained largely unchanged in the control groups. The levels of mRNAs of TERT and FST in brain, heart, kidney, liver, lung, and skeletal muscle from MCMV_TERT_ or MCMV_FST_ mice were 1.9 to 7.8 fold greater than WT treated controls in all tested organs (Fig. 2E, Supplementary Materials). The variations of the mRNA levels of TERT or FST in different tissues may be due to the different tropism of CMV and the post-transcriptional modification of TERT and FST. The relative telomere length in heart, liver, kidney, brain, lung, and muscle in 24-month-old MCMV_TERT_ treated mice was 6-fold greater than in control mice of the same age, and only ~8% shorter than an 8-month-old control (Fig. 2F) (25, 26, 39). (40). Antemortem daily observations did not reveal defects such as paralysis, body dysfunctions, or blindness. Visual histological analyses (not shown) revealed no malignancy or gross pathologies of the brain, liver, kidney, heart, muscle, bone, and lungs, in concordance with previous studies (9).

The relative telomere length in heart, liver, kidney, brain, lung, and muscle in 24-month-old MCMV_TERT_ treated mice was 6-fold longer than in control mice of the same age and only ~8% shorter than an 8-month-old control (Fig. 2F) (25, 26, 39). (40). Antemortem daily observations did not reveal defects such as paralysis, body dysfunctions, or blindness. Visual histological analyses (not shown) revealed no malignancy or gross pathologies of the brain, liver, kidney, heart, muscle, bone, and lungs, in concordance with previous studies (9).

### Hair and weight loss prevention

Bodyweight peaked at 23-months for all treatment groups except for MCMV_FST_ mice, whose weights continued to increase until 27-months and were ~33% heavier than the age-matched mock and WT controls. MCMV_TERT_ treatment also showed less weight loss over time compared to the mock and WT groups (Fig. 3B). Administration of MCMV_TERT_ and MCMV_FST_ was interrupted after mice reached 29 months of age when all mice in the control groups died (Fig. 3B, red arrow) but was resumed at 32 months (Fig. 3B, green arrow). When the treatment was stopped, the weights of MCMV_TERT_ and MCMV_FST_ groups declined, but the rate of weight loss decreased immediately upon therapy re-initiation. Future studies would be of interest to determine whether an uninterrupted monthly administration has a different outcome in longevity extension.

**Fig. 3.**
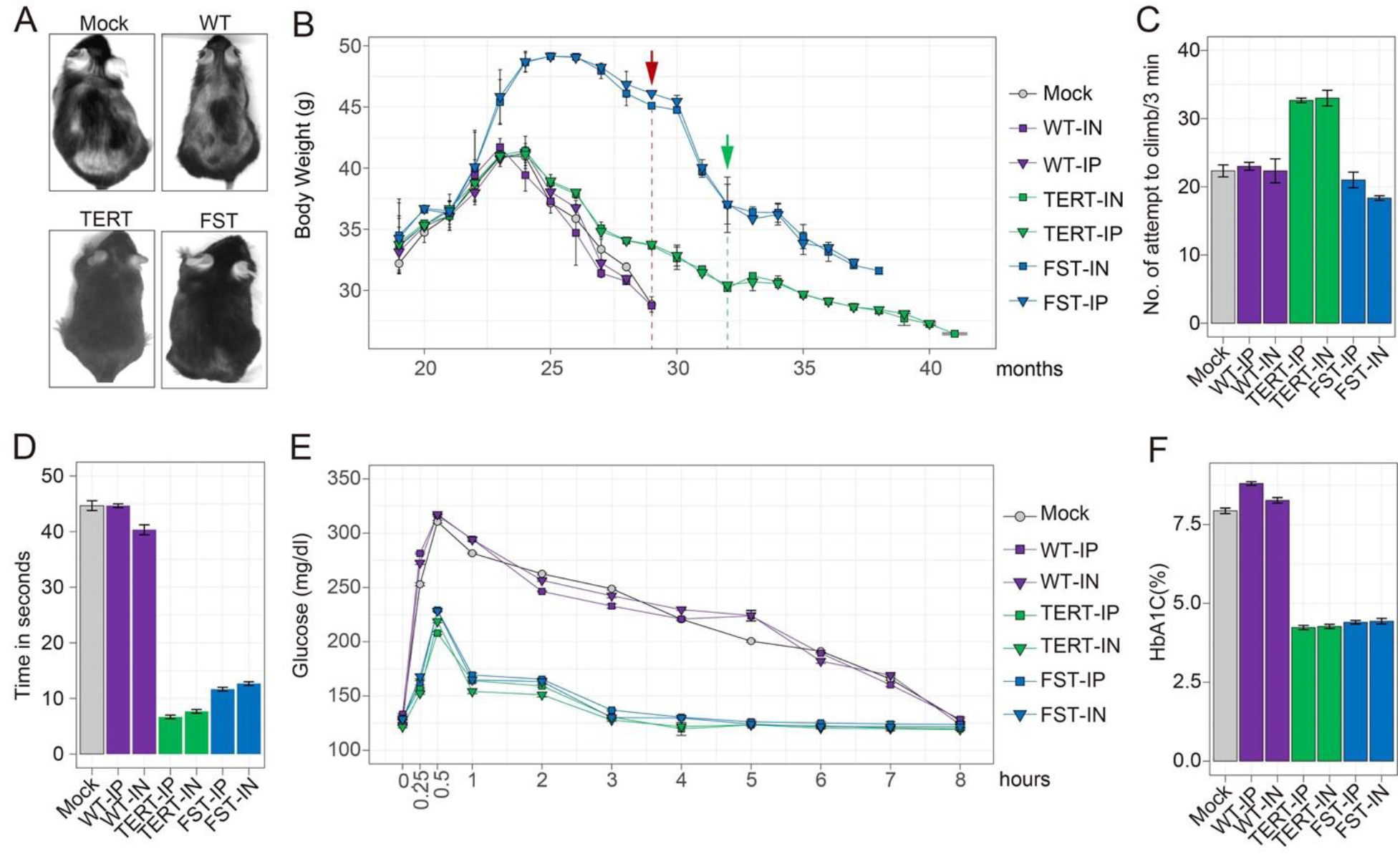
MCMV_TERT_ and MCMV_FST_ dramatically improved physical and physiological conditions. (**A**) Hair and body appearance after eight months of treatment. (**B**) Bi-weekly bodyweight averages of surviving mice in each group. Treatment interruption (red arrow) and reinitiation (green arrow). (**C**) Average number of climbing attempts in 3 minutes. (**D**) Beam crossing average execution time. (**E**) Glucose tolerance test. (**F**) HbA1c levels in Mock, WT, MCMV_TERT_, and MCMV_FST_ treated mice. Error bars represent the standard deviations.

### Improved activity and motor coordination

MCMV_TERT_ treated animals were ~40% more active than control mice in attempts to escape in a beaker test (30). Although MCMV_FST_ treated mice were bulkier and could not outperform the control mice in climbing attempts, they paced faster on the bottom of the beaker (Fig. 3C). Additionally, mice treated with MCMV_TERT_ or MCMV_FST_ completed a beam-crossing coordination test (31) in ~7.5s and 12.5s, respectively, as opposed to the controls (~43s), demonstrating superior coordination (Fig. 3D).

### Increased glucose tolerance

Glucose tolerance is known to decrease with aging. Here, we used a glucose tolerance test in fasted mice from each treatment group (Fig. 3E) (41). The average peak glucose concentration was ~33% lower for TERT and ~28% lower for FST treatments than for controls. Moreover, blood sugar levels reached baseline one hour post-administration in MCMV_TERT_ and MCMV_FST_ treated mice, in contrast to ~7 hours for control mice. In addition, the level of glycated hemoglobin (A1C) in treated mice was 4.5% and 4.7%, versus mock (7.9%) or WT (8.8%) (Fig. 3F). TERT and FST treatments were equally effective in blood glucose processing.

### Mitochondrial integrity in muscle

Mitochondria provide the essential metabolic support for an organism, and therefore play a central role in both lifespan determination and cardiovascular aging (42, 43). Here, we sacrificed 24-month-old mice from Mock, WT, MCMV_TERT_ and MCMV_FST_ treated groups, and sectioned heart and skeletal muscle tissues in order to examine subcellular structures of cardiomyocytes and skeletal muscle cells by electron microscopy. The number of mitochondria with connected cristae and mitochondrial area in the aged cardiomyocyte (Fig. 4A), as well as within cells of skeletal muscle (Fig. 4B) of MCMV_TERT_ or MCMV_FST_ treated mice were comparable to 6-month-old mouse controls, and substantially better than in age-matched control mouse tissues. These results suggest that MCMV_TERT_ and MCMV_FST_ preserved mitochondrial structure and sustained mitochondrial biogenesis.

**Fig. 4.**
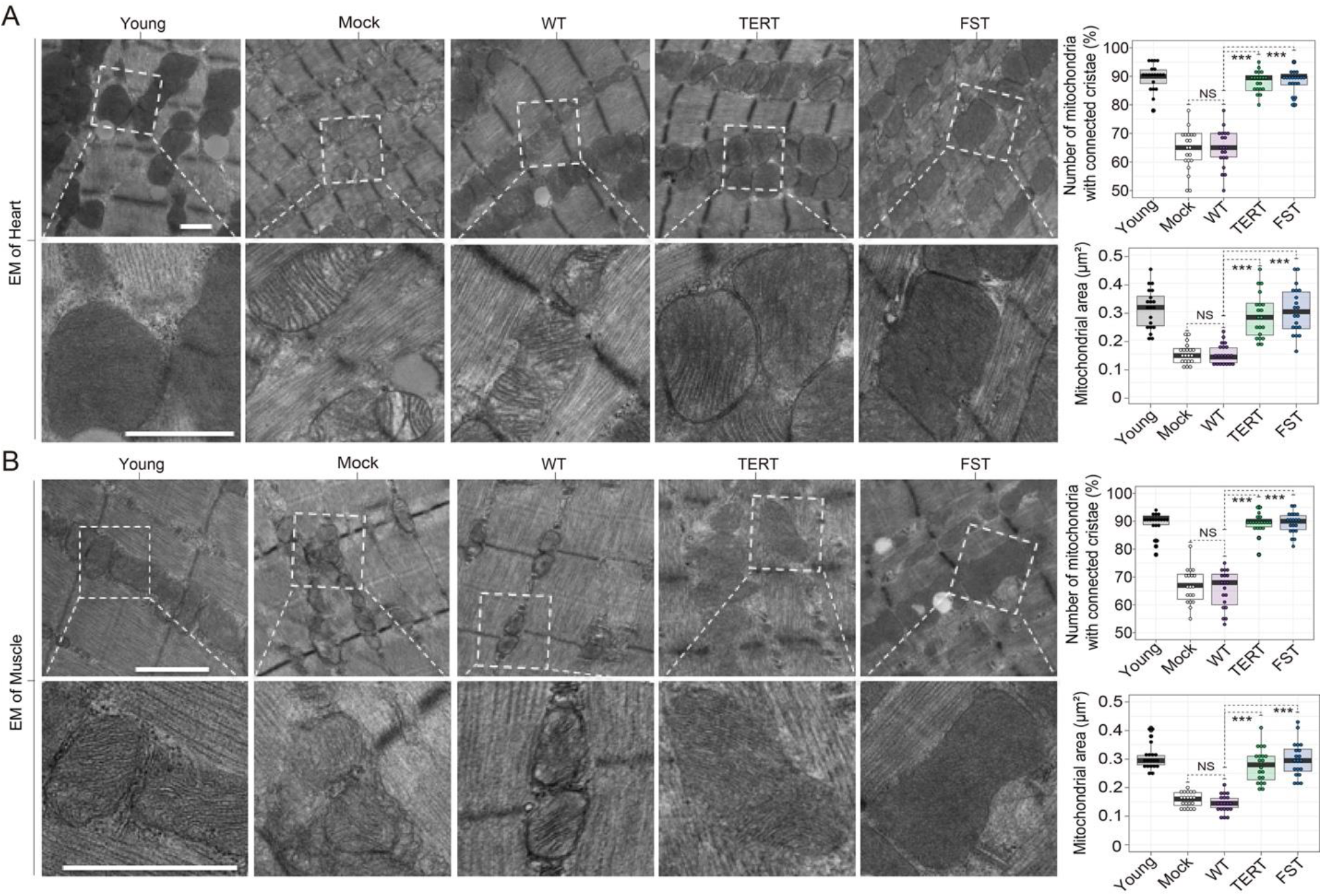
MCMV_TERT_ and MCMV_FST_ prevent mitochondrial deterioration in mice. (**A**) and (**B**) Representative EM images from the heart and skeletal muscle of untreated young mice and 24-month-old mice treated with Mock, WT, MCMV_TERT_, and MCMV_FST_ (IN groups are shown, scale bar=500nm). Quantitative analyses of the number of mitochondria with connected cristae and mitochondria area are on the right of A and B. *** *P*<0.001.

## DISCUSSION

Aging is often accompanied by the development of chronic conditions. The socioeconomic burden imposed by the diseases of aging could be lessened by maintaining a healthy aged population. We explored here a novel approach to achieve healthy aging, and show for the first time that CMV can be used as monthly inhaled gene therapy for delivering the exogenous genes TERT or FST safely and effectively. Interestingly, CMV therapy was equally effective regardless of route of inoculation, despite the fact that the mechanism of dissemination differs, which suggests that expression of the therapeutic load might not be substantially affected by the vector’s interaction with the immune system. Our results are congruent with other studies, which demonstrated that the olfactory route is a preferred natural route of CMV entry in murine models (37). Because herpesviruses are ubiquitous and acquired early, it has been proposed that a mutualistic relationship has developed between the co-evolving host and virus, where the latter even offers some immunomodulatory and homeostatic advantages to the hosts when in equilibrium (44). CMV’s large genome accommodates nearly 75% dispensable genes, many involved in immune evasion mechanisms that protect it from aggressive viral clearance responses regardless of route of entry. Olfactory infection spreads through dendritic cells, which migrate to lymph nodes and then extravasate into the bloodstream, whereas IP inoculation is expected to engage a wider range of myeloid progenitor cells, which in turn dictates viral dissemination and immune response outcomes (38).

FST treated mice showed an increase in body mass, as expected based on previous publications (45). We confirmed visually that skeletal muscle mass was indeed larger than that of controls at necropsy (data not shown). The increased robustness may explain the improved motor control in executing the beam test. However, it was unexpected that CMV-based FST gene therapy alone would increase longevity to the extent observed. Although it is known that FST has a concentration-dependent inhibiting effect on the myostatin-driven rate of muscle breakdown, which contributes to increasing frailty in aging individuals, the overall effect of increasing longevity warrants further inquiry. We anticipate that sarcopenia, muscular dystrophy, or even special circumstances causing muscle atrophy, such as low gravity exposure during space travel (46)., could be mitigated with a CMV-based FST gene delivery method.

Another surprising finding was the equivalent effectiveness of both treatment regimens in blood glucose control, because the cellular mechanisms activated by TERT and FST which ultimately result in glucose control are different. FST has a systemic role in upregulating factors controlling mitochondrial biogenesis, energy metabolism, cellular respiration and thermogenesis, inducing browning of white adipose tissue (47). The FST interference with the TGF-beta signaling pathway resulted in the efficient regulation of glucose homeostasis we observed. On the other hand, TERT seems to act at the level of pancreatic beta cells by upregulating insulin secretion rather than effecting glucose uptake (48). Nonetheless, telomerase is known to interact with various cellular inflammatory pathways to reduce oxidative stress, and has been detected in mitochondria, where it protects mitochondria from oxidative damage, which explains the systemic benefits and increased longevity (49). Furthermore, FST and TERT have shown positive effects in neurological diseases, and the fact that our treatment showed that brain FST and TERT levels increase significantly over baseline supports its use for treatment of these conditions (50, 51). It would be of great interest to understand the compounded effect the two therapies might have when delivered simultaneously.

Finally, our therapeutic regimen appeared to require monthly administration in order to have continuous effects, which may be advantageous when treatment indications do not require permanent expression of therapeutic load, but rather episodic or during specific circumstances, to achieve a reduced risk of long-term consequences in case of adverse reactions, should any occur.

In summary, our study justifies further efforts to investigate the use of CMV TERT and FST vectors against aging-related chronic inflammatory conditions, type 2 diabetes, sarcopenia, dementia, lung, kidney, and heart diseases responsible for decreased quality of life and premature death.

## Abbreviations

CMV: cytomegalovirus
DPI: days post-Inoculation and days post-Infection
FST: follistatin
IVIS: In Vivo Imaging System
MCMV: mouse cytomegalovirus
NCD: Noncommunicable diseases
PFU: plaque-forming-unit
TERT: telomerase reverse transcriptase
WHO: World Health Organization

## Acknowledgements

This work was fully funded by BioViva USA Inc, Delaware.

We thank Drs. Brian Kennedy, Shimon Meshi Zahav and Vivian Bellofatto for reviewing this study.

## Author contribution

Conceptualization: HZ, EP, AS; Methodology: HZ, EP, DKJ, GC, DK, QT, MT, JS, RC, SY; Investigation: HZ, EP, DKJ, QT, RC, MT, SY; Data analysis: HZ, EP, DKJ, DK, QT, MT, SY. Funding acquisition: HZ, EP; Writing – original draft: HZ, EP, AS, DKJ, SY, QT, MT; Writing – review & editing: GC, QT, AF.

## Data and material availability

The data are available in the main text or supplementary material files.

Correspondence and requests for materials should be addressed to H.Z.

---

**Supplementary table 1.**

| Groups | Virus | Number of Mice | Route of administration | Last mice died (Weeks) | Mice died (months) | Median age (Months) |
| --- | --- | --- | --- | --- | --- | --- |
| Group 1 | Mock | 8 | IP | 29.5 | 24.0, 25.0, 26.2, 26.4, 27.0, 27.2, 28.1, 29.5 | 26.7 |
| Group 2 | WT | 8 | IP | 28.1 | 23.5, 24.9, 26.1, 26.2, 26.7, 27.8, 28.0, 28.1 | 26.5 |
| Group 3 | WT | 8 | IN | 127.1 | 23.5, 24.6, 26.1, 26.2, 26.6, 26.8, 28.1, 29.5 | 26.4 |
| Group 4 | MCMV<sub>TERT</sub> | 8 | IP | 176.9 | 34.1, 34.3, 36.3, 37.2, 38.3, 38.4, 39.1, 41.2 | 37.8 |
| Group 5 | MCMV<sub>TERT</sub> | 8 | IN | 174.9 | 34.3, 35.2, 35.7, 36.4, 38.1, 38.4, 38.7, 40.5 | 37.3 |
| Group 6 | MCMV<sub>FST</sub> | 8 | IP | 161.9 | 31.5, 32.5, 33.8, 34.0, 35.9, 36.7, 36.8, 37.6 | 35.0 |
| Group 7 | MCMV<sub>FST</sub> | 8 | IN | 164.3 | 31.2, 32.1, 33.9, 34.9, 35.7, 36.9, 37.5, 38.0 | 35.3 |

